# *Cenchrus purpureus* and *Cenchrus americanus* repeatome provide chromosomal markers to distinguish subgenomes

**DOI:** 10.1101/2025.05.10.652685

**Authors:** Alex Junior Aparecido Silvestrini, Magdalena Vaio, Karla Carvalho Azevedo, Giovana Augusta Torres

## Abstract

*Cenchrus* L. is an important genus within the Poaceae family, comprising several species of high agronomic significance, such as *Cenchrus purpureus* and *Cenchrus americanus*, for production of forage and grains, respectively. *Cenchrus americanus* is a diploid species (2n = 2x = 14, AA genome), while *Cenchrus purpureus* is an allotetraploid (2n = 4x = 28, A’A’BB genome). The A’ subgenome is believed to be homeologous to and possibly derived from the A subgenome, while the origin of the B subgenome remains undefined. Despite their distinct subgenomic compositions, both species exhibit a high level of genome homology. The objective of the present work was the *in silico* characterization and comparative analysis of the repetitive fraction of the genomes of *Cenchrus purpureus* and *Cenchrus americanus* using genome skimming and a graph-based clustering approach, as well as the *in situ* hybridization of specific satellite DNA clusters into the chromosomes of both species. The repetitive fraction of the genome of *C. purpureus* and *C. americanus* corresponds to 52.23% and 76.82%, respectively. The most abundant repetitive elements in both species are the LTR retrotransposons. Satellite DNA sequences correspond to 2.55% and 4.17% of the genome of each species, respectively. Four new satellite sequences were identified as subgenome-specific sequences for both species, along with new centromeric variants. The ancestral relationship and the polyploidization-diploidization cycles played a fundamental role in the composition of their repetitive fraction. These cycles led *Cenchrus americanus*, a possible paleopolyploid, to a greater abundance of transposable elements compared to *Cenchrus purpureus*, a recent allopolyploid.

## Introduction

*Cenchrus* L. is a genus that belongs to the Poaceae family and has about 107 species distributed throughout the tropical and subtropical regions (Kew Gardens Science, 2020). The most recent taxonomic circumscription merged the previous genera *Cenchrus, Pennisetum* Rich. and *Odontelytrum* Hack under the single name *Cenchrus* (Chemisquy et al., 2010; Donadío et al., 2009). Several species of agronomic importance belong to *Cenchrus*, such as *Cenchrus purpureus* (Schumach.) (syn. *Pennisetum purpureum*), also known as Napier grass or Elephant grass, and *Cenchrus americanus* [(L.) R Brown, 1810] (syn. *Pennisetum glaucum*) known as Pearl millet. The first one is native to sub-Saharan Africa and due to its high dry mass yield, is an important forage species and a potential energy crop for tropical and subtropical regions of Asia, Africa and America. The second one is an important source of grains for human consumption and cultivated for fodder and fuel in arid and semi-arid regions of Asia and Africa (dos Reis et al., 2014; Sobrinho et al., 2005).

*Cenchrus* species are also of interest for genome evolution studies due to their complex genomic composition. *Cenchrus americanus* is a predominantly cross-pollinated diploid species (2n = 2x = 14, AA genome), while *Cenchrus purpureus* is an allotetraploid (2n = 4x = 28, A’A’BB genome), self-incompatible and obligate outcrossing species. The A’ subgenome is believed to be homeologous to and possibly derived from the A subgenome, while the origin of the B subgenome remains undefined. Both species exhibit a high level of genome homology (dos Reis et al., 2014; Paudel et al., 2018; Yan et al., 2021; Zhang et al., 2020) despite their distinct subgenomic compositions, reflecting their close phylogenetic relationship.

There are reports that *Cenchrus americanus* is a putative paleopolyploid, which, after diploidization processes, behaves as a diploid (Techio et al., 2006). In turn *Cenchrus purpureus* underwent at least three cycles of polyploidization events (Martel et al., 2004), with the most recent one about 15 MYA, coinciding with the origin of allopolyploidy. Some authors propose that the divergence of *Cenchrus purpureus* from *Cenchrus americanus* occurred at approximately 3.22 MYA (Yan et al., 2021), while others suggest it happened around 20 MYA (Zhang et al., 2022).

The genome sizes of *Cenchrus purpureus* and *Cenchrus americanus* are about 2.13-2.22 Gb and 1.85-2.32 Gb, respectively (Salson et al., 2023), and both species have complete genome assembly reports (Salson et al., 2023; Varshney et al., 2017; Yan et al., 2021; Zhang et al., 2022). As expected based on genome sizes, both genomes are enriched in repetitive DNA sequences.

Low-coverage genome sequencing, also known as genome skimming techniques (Dodsworth, 2015), combined with specific pipelines for studying genomic repetitive DNA (Novák et al., 2020), has been of great value for germplasm characterization and genome evolutionary studies (Gaiero et al., 2019; González et al., 2020; Heitkam et al., 2020; Sader et al., 2021). In this work, we conducted the first detailed and comparative description of the repetitive genome fraction of *Cenchrus purpureus* and *Cenchrus americanus*, aiming to better understand the genomic differentiation between them, as well as seeking evidences that could better explain the origin and evolution of the B subgenome.

## Results

### *Cenchrus purpureus* and *Cenchrus americanus* repetitive genome fraction

Comparative clustering analysis of the 1,060,996 reads from the *Cenchrus purpureus* (Cp) genome, identified 759,317 reads as repetitive sequences grouped into 75,634 clusters, while 301,679 were classified as singlets (Figure 1A). Those 759,317 reads after filtering by mitochondrial and chloroplast reads corresponded to approximately 52.23% of the genome (Figure 2, Table 1). From *Cenchrus americanus* (Ca) 1,313,846 reads were analyzed, of which 1,144,939 were allocated as repetitive sequences and 168,907 as singlets (Figure 1A). Ca repetitive sequences were grouped into 61,975 clusters, and the repetitive fraction of its genome corresponds to approximately 76.82% (Figure 2, Table 1). The 304 most abundant clusters, representing at least 0.01% of the genome, were analyzed and manually checked. Comparing the similarity between the analyzed clusters in both species, 2.63% of them are exclusive to the Cp genome, while 0.65% are exclusive to the Ca genome (Figure 1B), reflecting their genomic homology.

**Table 1.**
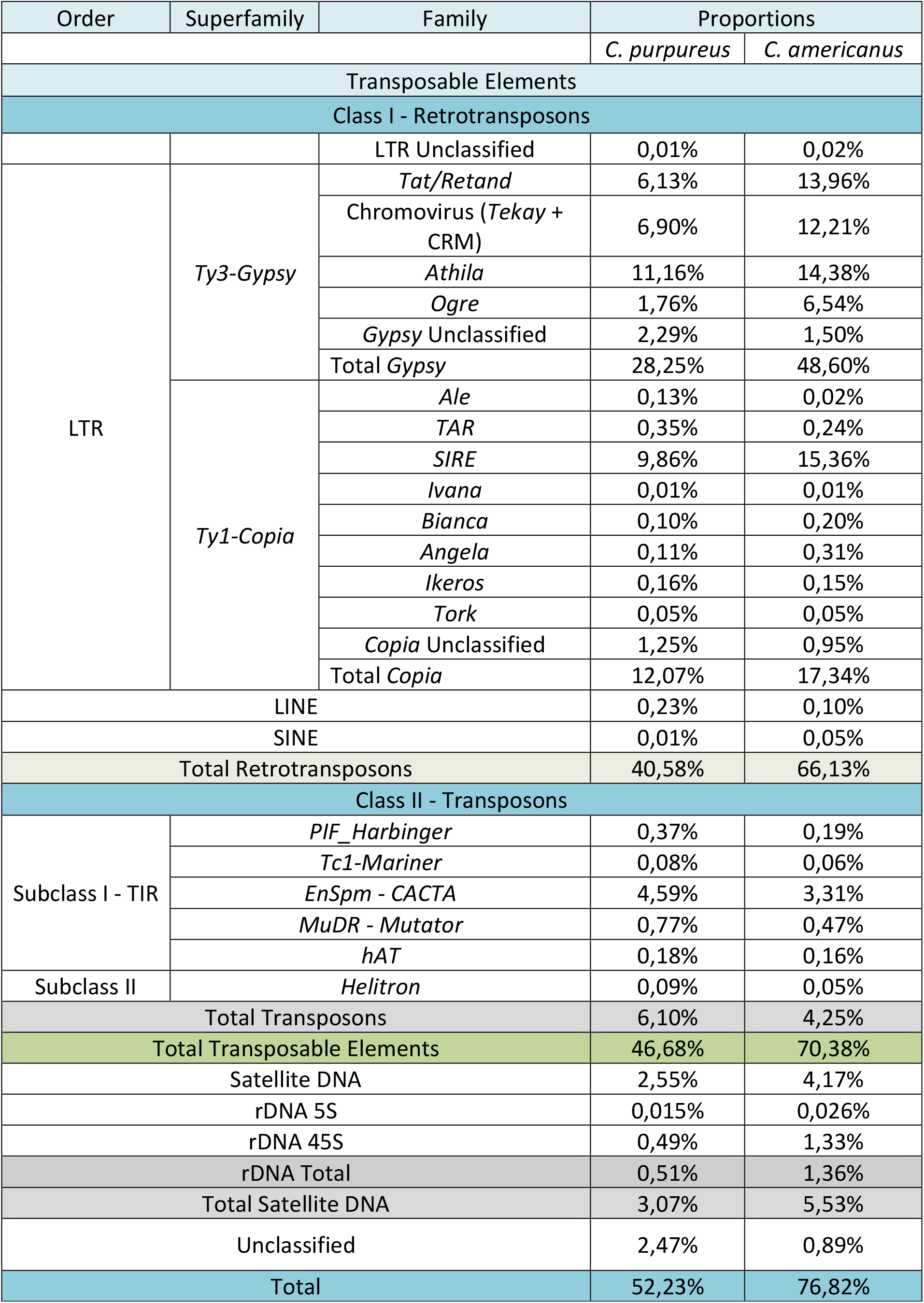
*Cenchrus purpureus* and *Cenchrus americanus* repetitive genome composition.

**Figure 1.**
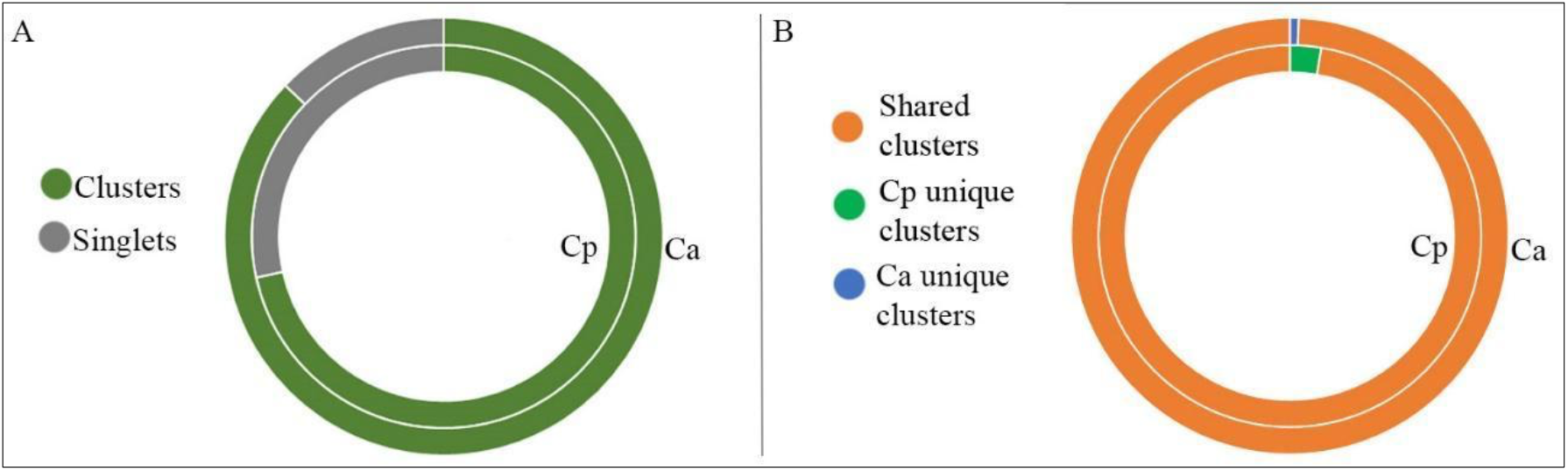
Comparative cluster summary of the analyzed *reads* by the RepeatExplorer platform in the genomes of *Cenchrus purpureus* and *Cenchrus americanus*. (A) Proportion of identified clusters and singlets from the RepeatExplorer*2* analysis. (B) Proportion of shared and exclusive clusters between *Cenchrus purpureus* (Cp) *and Cenchrus americanus* (Ca).

**Figure 2.**
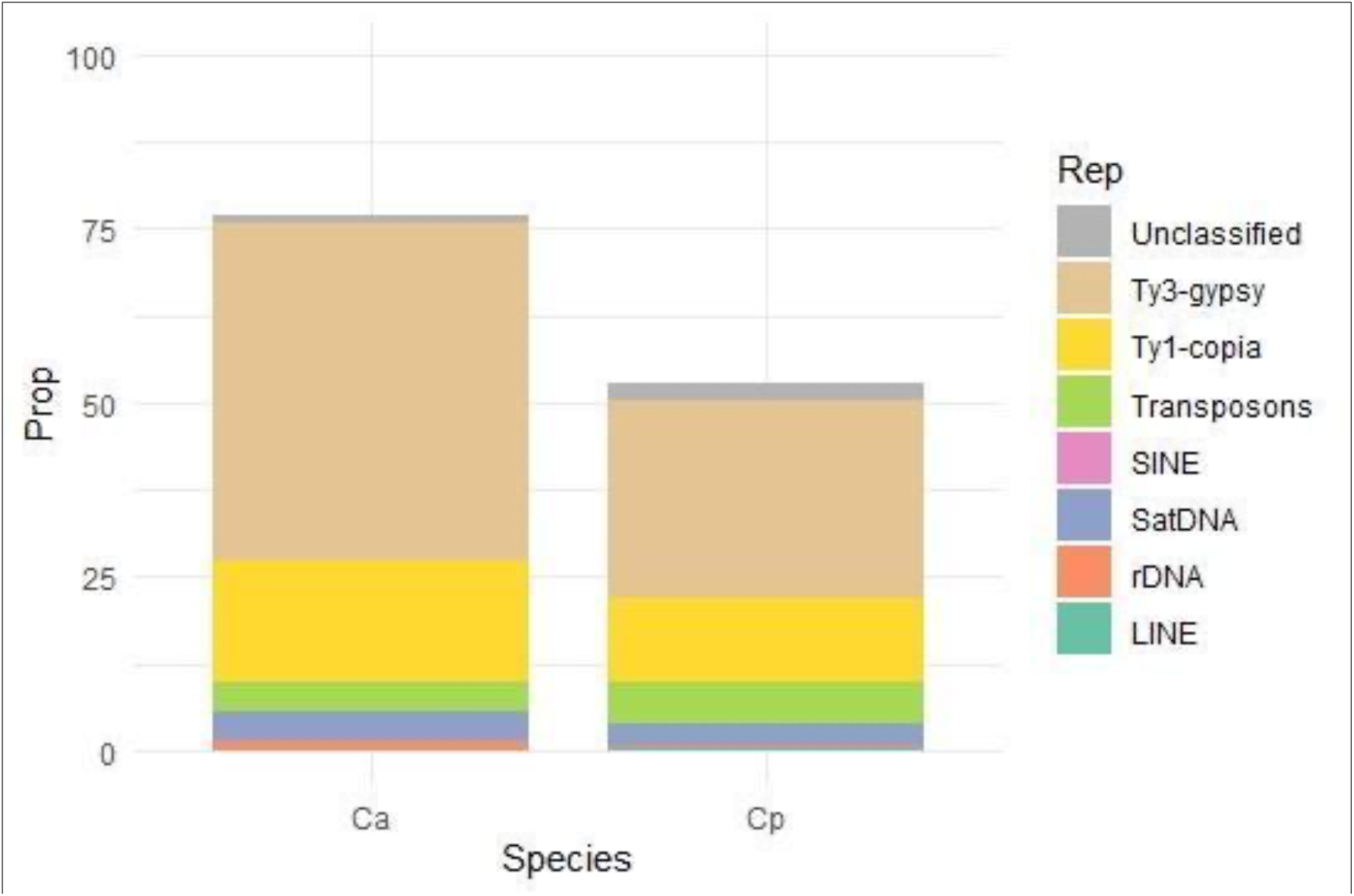
Repetitive fraction proportions in the genome of *Cenchrus purpureus (Cp)* and *Cenchrus americanus (Ca)*.

The retroelements are the most abundant class of repetitive sequences in both species, and the LTR order represents the highest fractions. *Ty3-Gypsy* superfamily is the most representative in the comparative analysis, with 48.60% of the Ca genome and 28.25% of the Cp genome (Table 1, Figure 2). The most abundant family within *Ty3-Gypsy*, in both genomes, is Athila. The second most abundant superfamily is *Ty1-Copia*, which corresponds to 17.34% and 12.07% of the genomes of Ca and Cp, respectively. In this case, the most representative family in both species is SIRE, with 9.86% in Cp and 15.36% in Ca. Regarding transposons, the most representative family is *EnSpm-CACTA* in both species (Table 1, Supplementary Figure 1).

### Phylogeny of the retrotransposase (RT) domains in *Ty3-gypsy* families

Phylogenetic analyses of the retrotransposase domain of *Ty3-gypsy* families (Figure 3) from Ca, Cp and *Setaria italica* (*S*i) proteome fraction showed that Tat/Retand clusters have differentiated within families in some lineages for each species, and thus they are recovered in different clades. The same pattern was obtained for Chromovirus/CRM lineages in which just a few are species-specific clades, while most of them are shared among all three species. There are lineages of *Ogre* and *Athila* specific to both *Cenchrus* species, and none of them are shared with Si. Some of the Tekay lineages have also diverged and are specific for each species, mostly from Ca and Cp genomes.

**Figure 3.**
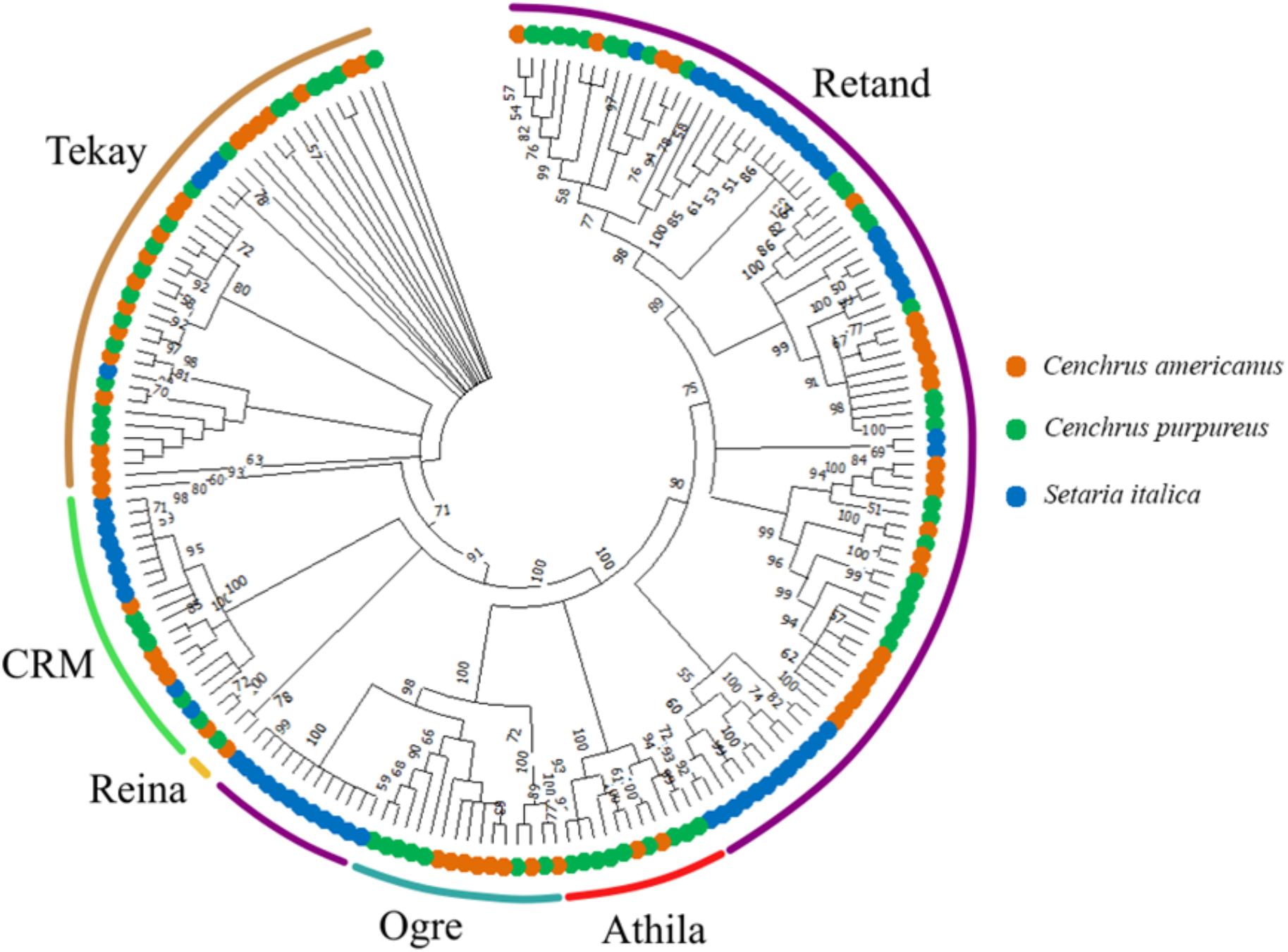
Phylogenetic relationships between the RT domains of Ty3-gypsy retroelement from *Cenchrus americanus, Cenchrus purpureus* and *Setaria italica* genomes. Bootstrap values are located at the nodes of each branch of the tree.

### Similarity of satellite DNA sequences from *Cenchrus purpureus* and *Cenchrus americanus* repeatome

According to the TAREAN analysis, the satellite DNA sequences correspond to 5.53% and 3.07% of the genome of Ca and Cp, respectively (Table 1, Figure 2). The Ca genome proportion occupied by sequences that encode for ribosomal gene 5S and 45S rDNA (18S, 5.8S and 26S) is 1.36%, while for Cp, it is 0.51%.

The TAREAN analysis of both species revealed fourteen satellite DNA sequences, ten from Cp and four from Ca genomes. Aiming to understand the evolutionary relationship between all these sequences, the monomers obtained from the TAREAN pipeline were aligned based on the RepeatMasker homology result. The monomers were classified into families and superfamilies (Table 2) based on their percentage of identity. In family 1 (F1; Suffix A), a total of four satellites were grouped. All F1 satellites have an average length between 137 and 156 bp and similar G/C content ranging from 41.7% to 44.2%. In family 2 (F2; Suffix B), two satellite DNA with 334 bp and 335 bp were identified. The G/C content of these satellites is 50% and 46.4%, respectively. Satellites identified in both species were grouped into the most specific family levels (F1 and F2), reflecting their genomic similarity. Superfamily 1 (SF1; Suffix Y) comprises six satellite DNA monomers with different lengths and G/C content. Superfamily 2 (SF2; Suffix Z) comprises two satellite DNA families.

**Table 2.** Satellite DNA sequences from *Cenchrus purpureus* and *Cenchrus americanus* genome. F/SF: Family/Super family. V: Number of variants. GC (%): GC content. Reads: Number of mapped reads from each satellite monomer. Cp: *Cenchrus purpureus*. Ca: *Cenchrus americanus*

| F/SF | DNASat | Length<br>(pb) | GC (%) | Abundance |  |  |  |
| --- | --- | --- | --- | --- | --- | --- | --- |
|  |  |  |  | Cp |  | Ca |  |
|  |  |  |  | *Reads* | Proportion<br>(%) | *Reads* | Proportion<br>(%) |
| F1 | CpaSat1A | 138 | 44,2 | 35629 | 2.912 | 28201 | 1.895 |
| F1 | CpurSat2A | 148 | 42,6 | 33216 | 2.715 | 25900 | 1.740 |
| SF1 | CpurSat3Y | 156 | 43,6 | 31812 | 2.600 | 25967 | 1.745 |
| F1 | CpurSat4A | 156 | 41,7 | 30664 | 2.506 | 25987 | 1.746 |
| SF1 | CpurSat5Y | 336 | 42,6 | 7534 | 0.616 | 0 | 0.000 |
| SF2 | CpurSat6Z | 154 | 51,3 | 7128 | 0.583 | 11419 | 0.767 |
| SF1 | CpurSat7Y | 372 | 43 | 6395 | 0.523 | 0 | 0.000 |
| F2 | CpurSat8B | 334 | 50 | 1887 | 0.154 | 11879 | 0.798 |
| SF2 | CpurSat9Z | 528 | 39,8 | 1069 | 0.087 | 0 | 0.000 |
| SF1 | CpurSat10Y | 364 | 43,7 | 272 | 0.022 | 0 | 0.000 |
| F1 | CameSat1A | 137 | 43,8 | 31857 | 2.604 | 26206 | 1.761 |
| F2 | CameSat2B | 335 | 46,3 | 1865 | 0.152 | 11887 | 0.799 |
| SF1 | CameSat3Y | 331 | 38,7 | 1435 | 0.117 | 336 | 0.023 |
| SF1 | CameSat4Y | 1314 | 48,1 | 18 | 0.001 | 220 | 0.015 |

All satellites were subjected to BlastN searches against the NCBI public database. Satellite sequence CpaSat1A, the most abundant (Table 3) and present in both genomes, presented hits against Gla’s cen, a known centromeric sequence previously isolated from Ca (Ishii et al., 2012). Homology levels were lower than 95%, thus CpaSat1A was classified as a different sequence belonging to the same family as Gla’s cen. Satellite CpurSat3Y also presented a hit against Gla’s cen, but with homology levels lower than 80%. Considering Gla’s cen was previously isolated from the Ca genome, and BlastN was not able to find identical hits (> 95% homology) in our TAREAN satellites, we performed a manual mapping of Gla’s cen against the pre-processed Ca reads. With that, we identified the centromeric sequence itself, with homology levels greater than 95%, and name this satellite as CameSat1A.

We mapped the consensus sequences of all the identified satellite DNA against the Illumina reads from both species, to assess their abundance (Figure 4). According to the mapping results, Cp presents all fourteen identified satellites, including those retrieved initially from the Ca genome. Family 1 (F1) had more than 33,000 and less than 35,000 reads mapped, indicating a large abundance and homogeneity of this specific family in the genomes. Family 2 (F2), on the other hand, presents reduced and heterogeneous abundance values, with more than 1,887 mapped reads and less than 11,887. Superfamily 1 (SF1) has elevated heterogeneity in the abundance pattern for each satellite sequence, ranging from 18 reads mapped on CameSat4Y to 31,812 reads on CpurSat3Y. SF1 is also heterogeneous, with satellites included in both species, while Superfamily 2 (SF2) presents only Cp satellite sequences (Figure 4).

**Figure 4.**
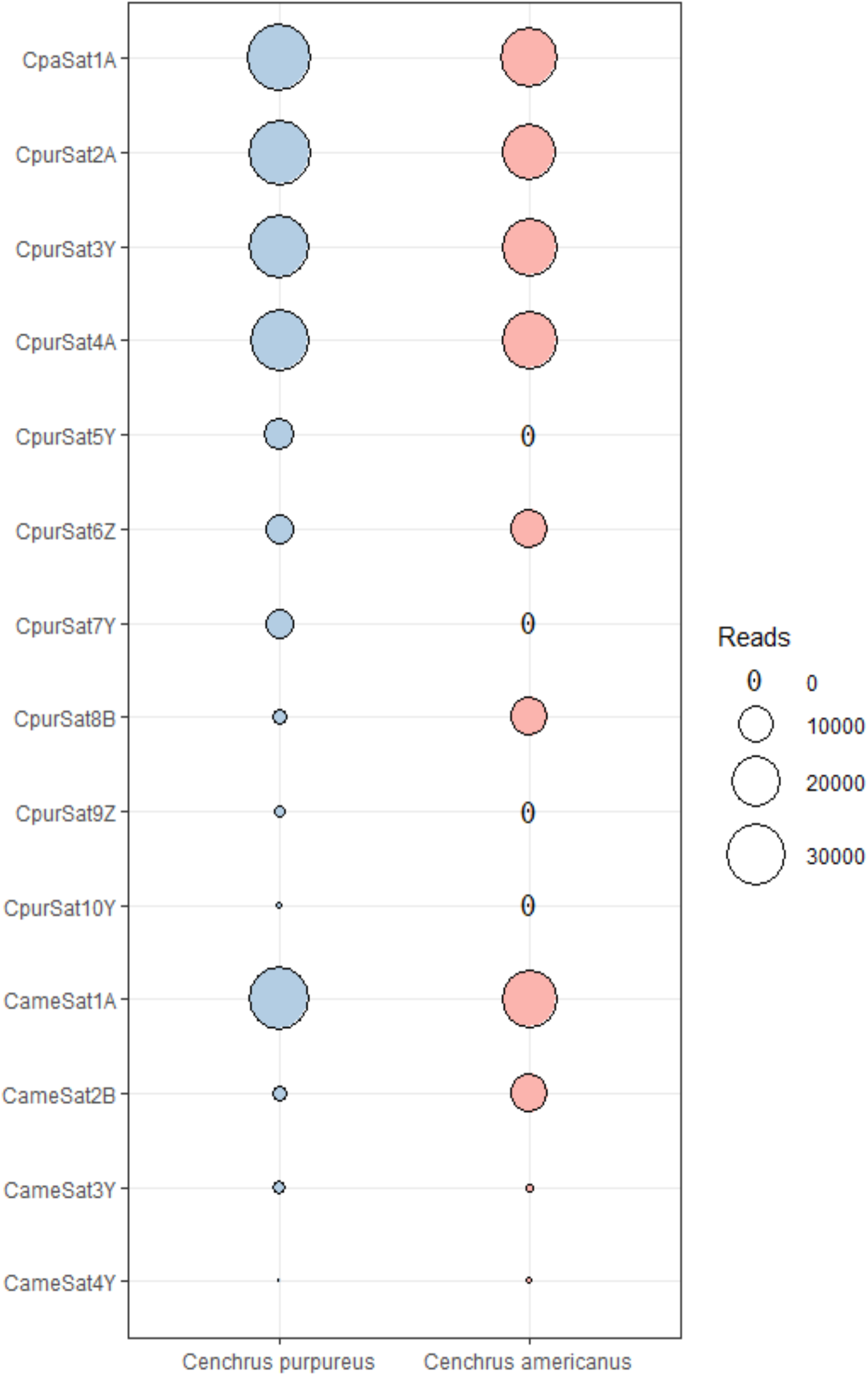
Bubble Chart showing the abundance of *reads* mapped from each satellite DNA monomer from *Cenchrus purpureus* and *Cenchrus americanus* genomes. CpurSat: Satellites originally extracted from the genome of *Cenchrus purpureus*. CameSat: Satellites originally extracted from *Cenchrus americanus*. CpaSat1A: Satellite that has variants in both genomes.

### FISH and whole genome mapping of *Cenchrus purpureus* and *Cenchrus americanus* satellite sequences

The chromosome-level genome assembly of Cp and Ca (Varshney et al., 2017; Yan et al., 2021) were used to map and define the presence or absence of specific satellite DNA families in the A, A’ and B subgenomes (Figure 5). Out of the fourteen described satellite sequences, thirteen were successfully mapped. Only CpurSat10Y did not yield any hits against the assembled genomes. Absent in the Ca genome, CpurSat10Y is present in low proportions in Cp short reads (Table 2, Figure 4), which may explain its absence in the genome assembly data.

**Figure 5.**
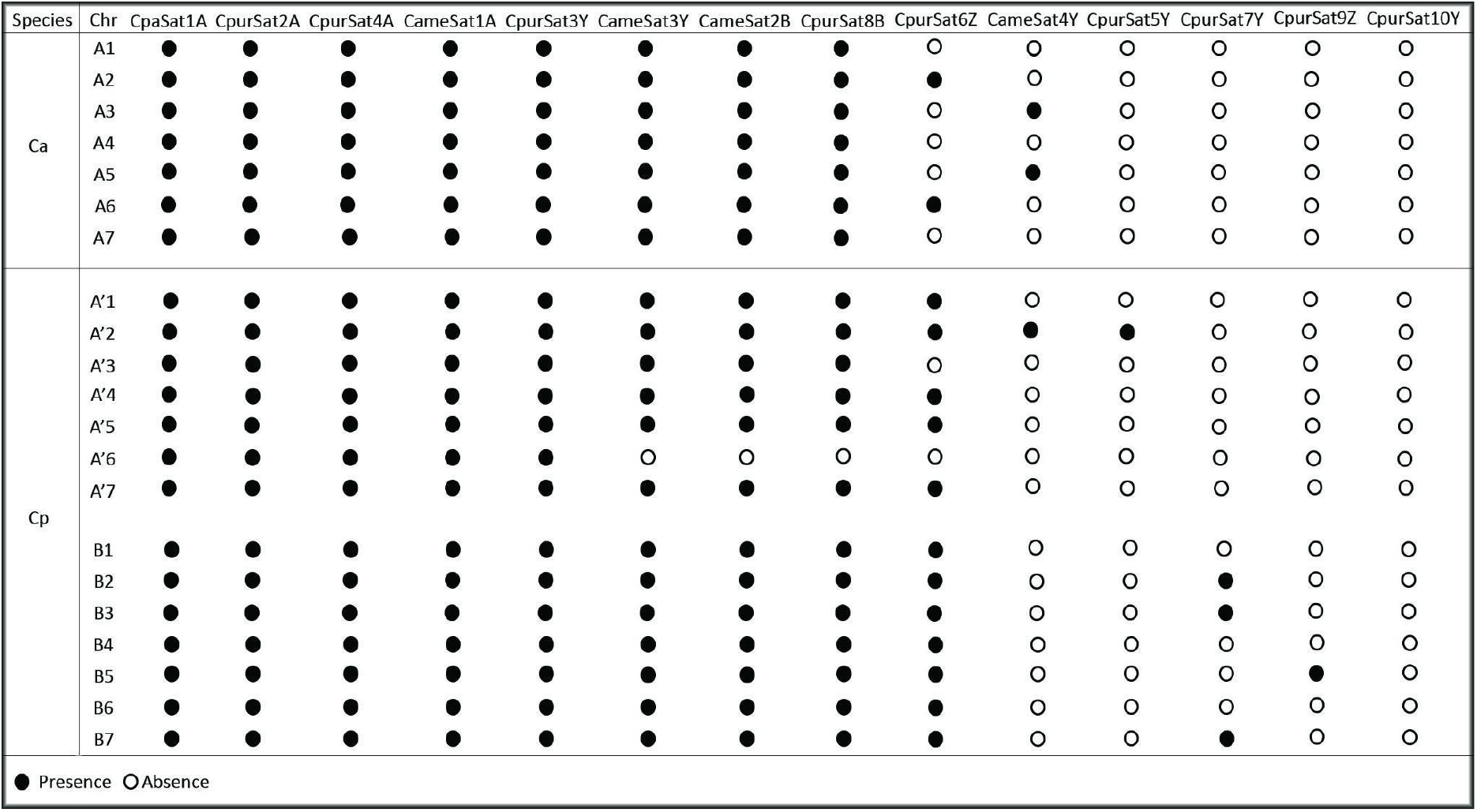
Genome assembly mapping presence (Black fill dot) and absence (White fill dot) of the identified satellite DNA monomers on each chromosome of *Cenchrus americanus* and *Cenchrus purpureus*. Ca: *Cenchrus americanus*. Cp: *Cenchus purpureus*. Chr chromosome number.

Five satellites (CpaSat1A, CpurSat2A, CpurSat3Y, CpurSat4A and CameSat1A), moslty belonging to F1, were mapped to all chromosomes of both species. Only three satellites and all belonging to Superfamily levels (CpurSat5Y, CpurSat7Y and CpurSat9Z) are species-specific, reflecting the high level of homology between both genomes. CameSat3Y (SF1), CameSat2B and CameSat8B (SF2) are absent only on chromosome A’6 of Cp. CpurSat6Z (SF2) is present on all chromosomes of the B subgenome but absent in four chromosomes of the A genome and two chromosomes of A’ subgenome. CpurSat4Y and CpurSat5Y (SF1) are exclusive to the A and A’ subgenomes. CpurSat7Y and CpurSat9Z (SF1) are found only on B subgenome chromosomes (Figure 5).

We performed fluorescent *in situ* hybridization (FISH) experiments using selected satellite sequences identified through the *in silico* mapping as potential subgenome markers (Figure 5): CameSat2B as A, A’ and B marker, CpurSat9Z and CpurSat7Y as B markers and CpurSat5Y as an A’ marker. We also used the likely new centromeric satellite variant of *Cenchrus*, CpaSat1A, to confirm its chromosomal location. As shown in Figure 6A, CpaSat1A is indeed a centromeric variant of the previously isolated satellite Gla’s cen. CpaSat1A hybridized to the proximal/centromeric regions in all chromosomes in both Cp and Ca, confirming our results.

**Figure 6.**
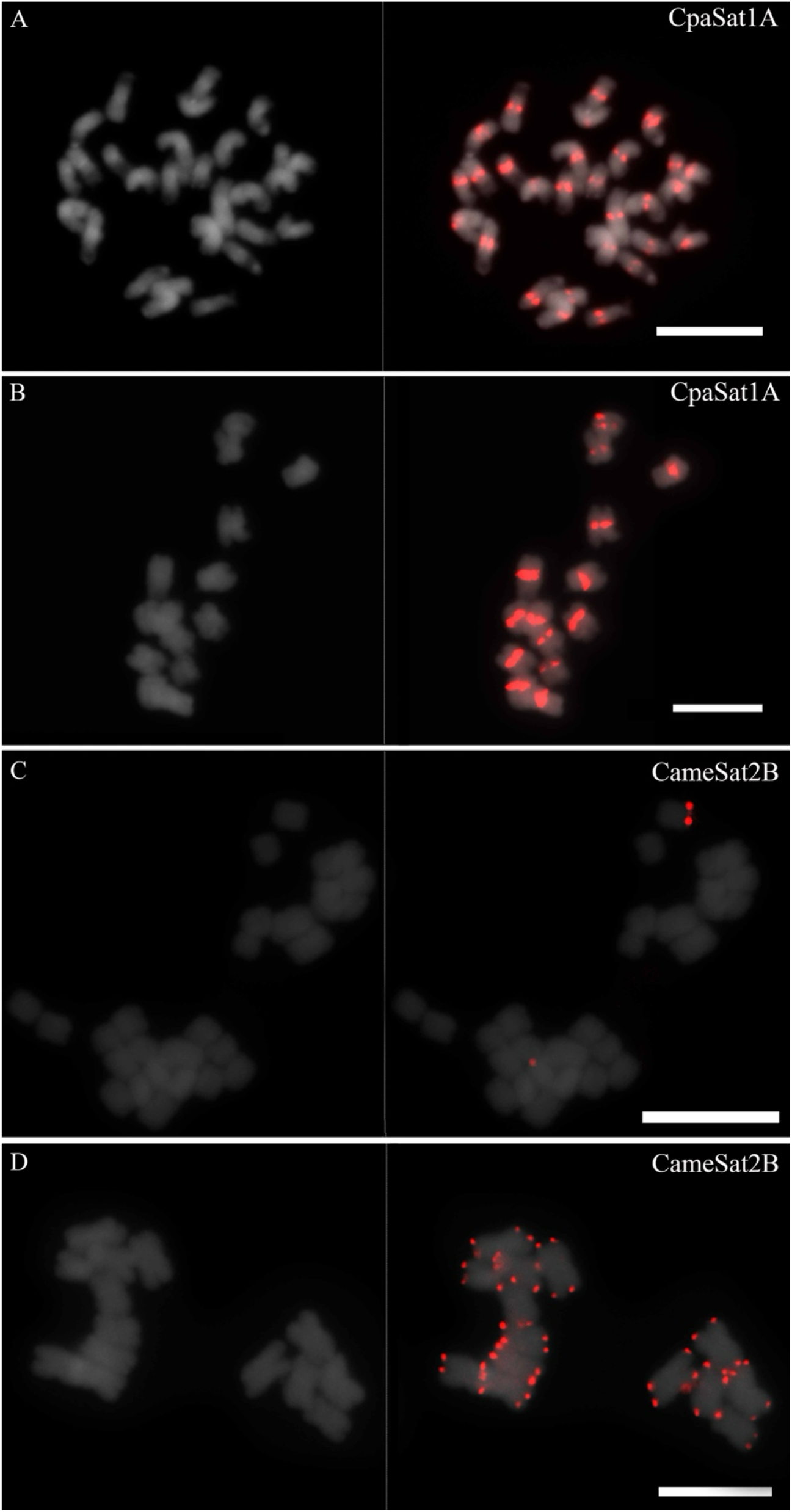
Fluorescent *in situ* hybridization mapping of two satellite sequences (CapSat1A and CameSat2B) on DAPI counterstained chromosomes of *Cenchrus purpureus* (A and C) and *Cenchrus americanus* (B and D). Bar = 10 µm

**Figure 7.**
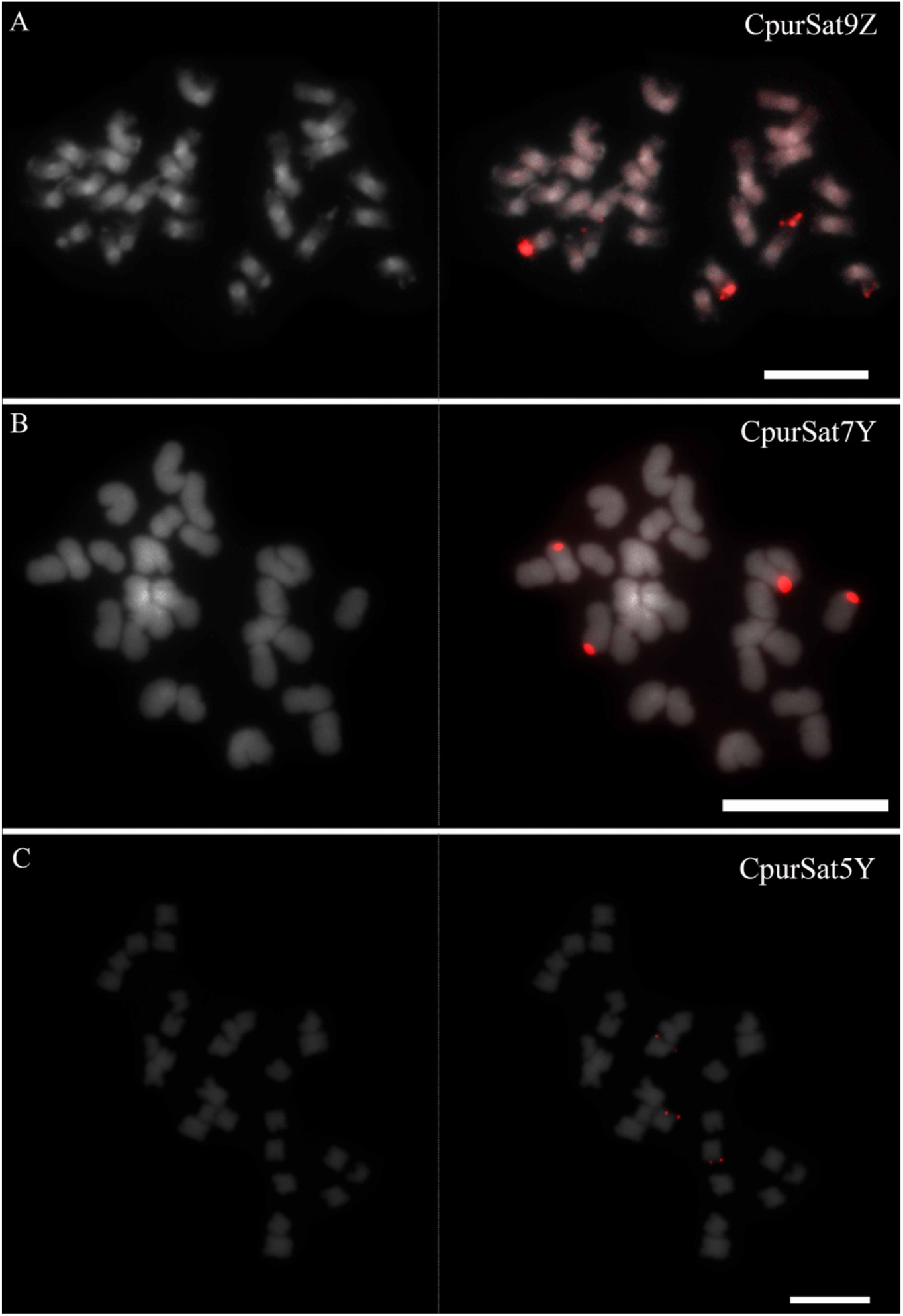
Fluorescent *in situ* hybridization mapping of three satellite sequences (CpurSat9Z, CpurSat7Y, CpurSat5Y) on DAPI counterstained chromosomes of *Cenchrus purpureus*. Bar = 10 µm.

None of CpurSat9Z, CpurSat7Y or CpurSat5Y hybridized to Ca (A subgenome) chromosomes, indicating that all three can be considered markers for the A’ or B subgenome (Figure 6B). These results corroborate the genome assembly mapping (Figure 5) and reveal that CpurSat9Z and CpurSat7Y are B specific markers and CpurSat5Y is a A’ specific marker. CameSat2B hybridized to subtelomeric regions of all chromosomes in Ca and to only one chromosome in Cp (Figure 6A). This sequence represents a newly described subtelomeric sequence in Ca. The pattern of CameSat2B signals in Cp contradicts the *in silico* genome assembly mapping results (Figure 5). This can be explained either by the challenge of assembling perfect satellite monomers (Hu et al., 2024) or by insufficient number of repeats for FISH detection.

## Discussion

Advances in sequencing technologies have enabled the characterization of the repetitive genome fraction in non-model plant species. In the present work, we performed a complete characterization of the repetitive genome fraction using genome skimming data from two grass species of high importance for livestock, *Cenchrus purpureus* (2n = 4x = 28=A’A’BB), and human food, *Cenchrus americanus* (2n=2x=AA). Both species share an A genome common ancestor about 20 Mya (Salson et al., 2023). The *in silico* studies revealed high similarity among the three subgenomes and the presence of species and subgenome-specific satellite DNA families, allowing the identification of specific A, A’ and B chromosomes.

### Differences in the repeat composition of *Cenchrus americanus* and *Cenchrus purpureus* can be related to ancient WGD events

Plant genomes often contain large proportions of repetitive sequences, which can comprise up to 80% in relatively large genomes (Novák et al., 2014). The repetitive genome fractions of *Cenchrus purpureus* (52.23% - 2.22 Gb) and *Cenchrus americanus* (76.82% - 2.28 Gb) fall within the expected range for species with genome sizes between small and intermediate (1 .40 Gb – 13.71 Gb) (Soltis et al., 2003), and are consistent with genome assembly reports showing ∼66% and ∼77%, respectively (Salson et al., 2023; Varshney et al., 2017; Yan et al., 2021; Zhang et al., 2022).

*Cenchrus americanus*, a diploid species, has a relatively larger repetitive genome fraction, with its transposable element content being 1.6 times greater than that of *Cenchrus purpureus*, an allotetraploid. *Cenchrus americanus* is a putative paleopolyploid, which, after cycles of polyploidization-diploidization, behaves like a diploid species (Martel et al., 2004; Techio et al., 2006). In contrast, *Cenchrus purpureus* is a recent allopolyploid (dos Reis et al., 2014; Martel et al., 2004; Zhang et al., 2020) that originated through a hybridization event between the common ancestor of both species and an unknown ancestor with a genome B composition.

Processes that lead to numerical and/or structural chromosomal changes, as already described for *Cenchrus americanus* (dos Reis et al., 2014; Martel et al., 2004; Paudel et al., 2018; Varshney et al., 2017), can induce genomic stresses causing an exponential increase in the repetitive fraction and, consequently, in the genome size (Belyayev, 2014). This may explain the relatively larger genome of Ca with a higher abundance of transposable elements compared to Cp, since the first has undergone at least three polyploidization-diploidization cycles (Techio; Davide; Pereira, 2006). The differential amplification of specific retrotransposon lineages in Ca, as reported in the phylogenetic analyses involving *Ty3-gypsy* families, corroborate these findings, indicating a possible increase in the diversity of these elements in the Ca genome.

By comparing the exclusive and shared clusters between both species, we observed a high degree of similarity in the repetitive DNA sequences from Ca and Cp, which can be explained by the high level of similarity between all three subgenomes – A, A’ and B. This similarity has already been attested by genome synteny and genomic *in situ* hybridization (GISH) analysis (Dos Reis et el., 2014; Yan et al., 2020).

### Genome mapping and FISH using satellite DNA sequences allow the identification of putative new centromeric sequences and subgenome-level chromosomal markers in *Cenchrus purpureus* and *Cenchrus americanus*

*Cenchrus americanus* and *Cenchrus purpureus* present low abundances of satellite sequences (4.17 and 2.55%, respectively) compared to transposable elements (70.38 and 46.68%, respectively). These results are in line with those proposed for several other grasses such as rice, maize, and sorghum (Bilinski et al., 2017). Ca has a higher proportion of satellite sequences than Cp. This suggests a correlation between the loss of repetitive sequences, mainly satellite DNA sequences, and the event of allopolyploidization (Wang et al., 2018; Zhang et al., 2020).

Genome assembly mapping and FISH results show a new centromeric variant for *Cenchrus*. Considering the similarity between CameSat1A and CpaSat1A to *Gla’s cen*, a known centromeric satellite, as well as the CpaSat1A FISH mapping to the centromere, we propose that CameSat1A and CpaSat1A are centromeric sequences in Ca and Cp chromosomes. CpaSat1A can also be proposed as a variant (< 95%) of *Gla’s cen*, which was not previously reported.

The data from the subgenome-level mapping corroborate the results concerning the abundance and FISH of satellite DNA families. The satellites present in all chromosomes of both species (CpaSat1A, CpurSat2A, CpurSat3Y, CpurSat4A and CameSat1A) mainly represent those classified at the family level, more specifically those belonging to F1. These sequences are putatively present in the common ancestor between *Cenchrus americanus* and *Cenchrus purpureus* because they are present in the three subgenomes that compose both species – A, A’ and B (Figures 5 and 6). These data suggest a certain degree of similarity between the genome of the unknown B donor and the ancestors of both species, which is proposed to have an A composition (dos Reis et al., 2014).

Considering the homeology between A and A’, the presence of CameSat2B in all Ca chromosomes and only one Cp, and the challenge to assemble the repetitive fraction of larger genomes, we can propose CameSat2B as an A and A’ subgenome FISH marker (Figures 5 and 6A). Satellites differentiated at the super-family level, such as CpurSat9Z, CpurSat7Y and CpurSat5Y, are completely divergent from other identified satellite sequences in both species, thus classified at the super-family level. In the FISH results, those satellites can be used as subgenome markers in Cp, both for A’ and B subgenomes, since they only hybridize to Cp chromosomes.

The genomic homology between A and A’ and the degree of dissimilarity between these two subgenomes and the B subgenome as proposed by dos Reis et al., (2014) may also explain the presence of specific A/A’ and B satellites. A and A’ specific satellites are likely present in the A genome progenitor of both species, while B-specific satellites are present only in the unknown B donor. The identified subgenome-specific sequences can be used as robust markers for subgenome-chromosome identification in other *Cenchrus* species, to understand better how these subgenomes evolved and diverged.

## Methods

### Plant genotypes and Illumina Hiseq™ 4000 genome sequencing

Young leaves of *Cenchrus purpureus* cv. Pioneer (Cp) and *Cenchrus americanus* cv. BN2 (Ca) were used for DNA extraction using the DNeasy® Plant Mini Kit (Qiagen® Inc., Venlo, Netherlands). The sequencing library preparation was performed using the Nextera™ DNA Flex Library Prep kit with 2×300bp pair-ended fragments for both species. Genome sequencing was performed on the Illumina HiSeq™ 4000 platform at the René Rachou Institute – Fiocruz Minas, Belo Horizonte – MG - Brazil.

### Comparative analysis of whole genome repetitive fraction and transposable elements characterization

The characterization of the repetitive fraction was performed by a similarity-based clustering analyses using the RepeatExplorer2 pipeline (Novak et al., 2020) at the Elixir-Cerit server (https://repeatexplorer-elixir.cerit-sc.cz/) using default parameters. First, sequencing reads from both species were uploaded to the server and filtered by quality using a cut-off value of 10 in at least 95% of the base pairs. After discarding low-quality sequences, we performed a read sampling that was equivalent to the Cx (genome size per subgenome) of the two species, to equate the differences in the ploidy level between both species (Gaiero et al., 2019). Sequencing reads of each species were identified with a prefix: Cp for *Cenchrus purpureus* and Ca for *Cenchrus americanus*, concatenated and submitted to comparative clustering analyses following Novak et al., 2020. Clustering analyses were performed using default settings of RepeatExplorer2.

Automatic annotation of the repetitive elements present in both genomes was generated from the clustering analysis using the Viridiplantae database of protein domains for plant transposable elements – REXdb (Neumann et al., 2019). Unclassified transposable elements that represented above 0.01% of the genome were manually checked against the Giri Repbase (Bao et al., 2015) and classified according to their similarity to previously deposited repetitive elements. Clusters representing chloroplast and mitochondrial DNA were excluded before calculating repeat abundance for each species. The proportion of each type of repetitive sequence was calculated considering the number of reads for each cluster and the total reads included in the clustering run.

The most abundant transposable element family was selected for further phylogenetic analysis using its retrotransposase (RT) domain. The DANTE (Domain based ANotation of Transposable Elements) algorithm (Neumann et al., 2019; Novák et al., 2020) implemented in RepeatExplorer was used to search and extract the component amino acid sequences of the conserved retrotransposase domain of all the *Ty3-gypsy* retroelements of both species. Sequences of the retrotransposase (RT) protein domains from *Setaria italica* (Si) were used as an outgroup (Chemisquy et al., 2010). The amino acid sequences were aligned using the MUSCLE algorithm, and the phylogenetic relationships were calculated using the Neighbour-Joining method, including bootstrap values. All analyses were performed using MEGA-X software (Kumar et al., 2018).

### Detailed characterization, classification and *in silico* genome assembly mapping of satellite DNA families

The satellite characterization was performed using TAREAN (Tadem Repeat Analyzer), also implemented in RepeatExplorer (Novák et al., 2020). For this, we used each species’ pre-processed short reads, corresponding to approximately 0.1x their genome sizes. All clusters classified as satellite DNA for each species were further analyzed using Dot Plot graphs available from the EMBOSS package (Rice et al., 2000) to confirm their tandem arrangement. The satellite DNA monomers generated by TAREAN were subject to similarity searches using BLASTn at <https://blast.ncbi.nlm.nih.gov/Blast.cgi>.

The homology, relationship and identity among the satellite DNA monomers were analyzed using the Repeat Masker v4.05 software (SMIT et al., 2015) and the global sequence alignment algorithm present in MAFFT (Katoh & Standley, 2013). The satellite monomers were then classified into families and subfamilies according to their identity. Identity values greater than 95% among different monomers were considered as one variant of the same satellite DNA, values between 80% and 95% were classified as different variants belonging to the same satellite DNA family, and identity below 80% as belonging to the same satellite DNA superfamily (RUIZ-RUANO et al., 2016).

The satellite sequences were organized and enumerated according to their abundance related to their respective identified species. To differentiate between families and superfamilies, suffixes were assigned according to the initial letters of the Roman alphabet, A and B, for the family level and the final letters, Y and Z, for the superfamily level. The relative quantification of each satellite DNA family was performed through the mapping of all the obtained monomers against the raw Illumina reads from *Cenchrus americanus* and *Cenchrus purpureus* using the standard read mapping algorithm implemented on Geneious Prime 2021.0.3. Genome assembly (Varshney et al., 2017; Yan et al., 2020) mapping of satellite families was done using the standard genome mapping algorithm implemented on Geneious Prime 2021.0.3 (Kearse et al., 2012).

### Satellite sequences amplification, slides preparation and Fluorescent *in situ* Hybridization (FISH)

#### Plant Material and Chromosome Preparation

Young root tips of *Cenchrus americanus* and *Cenchrus purpureus* were harvested and pretreated with 89 µM cycloheximide for 2 hours at 24°C. Subsequently, the material was fixed in methanol:acetic acid (3:1, v/v) for 24 hours at room temperature and stored at −20°C until use. For chromosome preparation, root tips subjected to enzymatic digestion (0.7% cellulase Onozuka R10, 0.7% cellulase, 1% pectolyase, and 1% cytohelicase) at 37°C for 2 hours in a water bath. Chromosome spreads were prepared using the flame-drying method (Dong et al., 2000). To reduce cytoplasmic background, slides were pretreated with 10 µg/mL pepsin at 37°C for 30 minutes, washed in distilled water and 2× SSC buffer, followed by post-fixation in 4% formaldehyde for 10 minutes and dehydration through an ethanol series.

#### Fluorescence In Situ Hybridization (FISH)

Five satellite DNA probes (Supplementary Table 1) were amplified via PCR using genomic DNA extracted from their respective source species. Primer pairs were designed to target consensus sequences using the Primer3 tool with default parameters (Koressaar et al., 2018). Genomic DNA from *C. americanus* and *C. purpureus* was extracted from young leaves of greenhouse-grown plants following the CTAB protocol (Doyle and Doyle, 1987). DNA integrity was verified by 1% agarose gel electrophoresis, and concentrations were quantified using a NanoDrop™ spectrophotometer prior to probe labeling via nick translation with either biotin-14-dATP or digoxigenin-11-dUTP.

Chromosomes were denatured in 70% formamide at 80°C for 1 minute, followed by dehydration in an ethanol series. FISH procedures were performed according to Dong et al., 2000. The hybridization mixture (50-100 ng/µL of each double-stranded DNA probe, 50% formamide, 20% dextran sulfate, 20× SSC) was denatured at 95°C for 10 minutes and applied directly onto the pre-denatured chromosome slides. Hybridization was carried out at 37°C overnight. Detection was performed using anti-digoxigenin antibodies conjugated to rhodamine or anti-biotin antibodies conjugated to FITC. Slides were mounted in Vectashield® mounting medium containing DAPI (4′,6-diamidino-2-phenylindole). Fluorescent signals were analyzed using an epifluorescence microscope equipped with a cooled digital camera and specific filter sets: DAPI (excitation: 330–385 nm; emission: 420 nm), FITC (excitation: 490 nm; emission: 525 nm), and rhodamine (excitation: 400–410 nm; emission: 455 nm). Images were processed using Adobe Photoshop.

## Acknowledgments

We thank CAPES, FAPEMIG and CNPq for the project funding and Fiocruz - Rene Rachou Institute for the genomic sequencing.

## Notes

### Competing Interest Statement

The authors have declared no competing interest.

## References

Andrade-Vieira, L. F. (2010). Anslise Citomolecular em hibridos de campim-elefante e milhero. (Pennisetum sp. Schum ., Poaceae). Universidade Federal de Lavras.

Bilinski, P., Han, Y., Hufford, M. B., Lorant, A., Zhang, P., Estep, M. C., Jiang, J., & Ross-Ibarra, J. (2017). Genomic abundance is not predictive of tandem repeat localization in grass genomes. PLoS ONE, 12(6), 1–12. 10.1371/journal.pone.0177896

Chemisquy, M. A., Giussani, L. M., Scataglini, M. A., Kellogg, E. A., & Morrone, O. (2010). Phylogenetic studies favour the unification of Pennisetum, Cenchrus and Odontelytrum (Poaceae): A combined nuclear, plastid and morphological analysis, and nomenclatural combinations in Cenchrus. Annals of Botany, 106(1), 107–130. 10.1093/aob/mcq090

Dodsworth, S. (2015). Genome skimming for next-generation biodiversity analysis. Trends in Plant Science, 20(9), 525–527. 10.1016/j.tplants.2015.06.012

Donadío, S., Giussani, L. M., Kellogg, E. A., Zuolaga, F. O., & Morrone, O. (2009). A preliminary molecular phylogeny of Pennisetum and Cenchrus (Poaceae-Paniceae) based on the trnL-F, rpl16 chloroplast markers. Taxon, 58(2), 392–404.

dos Reis, G. B., Mesquita, A. T., Torres, G. A., Andrade-Vieira, L. F., Pereira A. Vander, & Davide, L. C. (2014). Genomic homeology between Pennisetum purpureum and Pennisetum glaucum (Poaceae). Comparative Cytogenetics, 8(3), 199–209. 10.3897/CompCytogen.v8i3.7732

Gaiero, P., Vaio, M., Peters, S. A., Schranz, M. E., de Jong, H., & Speranza, P. R. (2019). Comparative analysis of repetitive sequences among species from the potato and the tomato clades. Annals of Botany, 123(3), 521–532. 10.1093/aob/mcy186

González, M. L., Chiapella, J., Topalian, J., & Urdampilleta, J. D. (2020). Genomic differentiation of Deschampsia antarctica and D. cespitosa (Poaceae) based on satellite DNA. Botanical Journal of the Linnean Society, 1–16. 10.1093/botlinnean/boaa045

Heitkam, T., Weber, B., Walter, I., Liedtke, S., Ost, C., & Schmidt, T. (2020). Satellite DNA landscapes after allotetraploidisation of quinoa (Chenopodium quinoa) reveal unique A and B subgenomes. The Plant Journal, March, tpj.14705. 10.1111/tpj.14705

Hu, J., Wang, Z., Liang, F., Liu, S.-L., Ye, K., & Wang, D.-P. (2024). NextPolish2: A Repeat-aware Polishing Tool for Genomes Assembled Using HiFi Long Reads. Genomics, Proteomics & Bioinformatics, 22(1). 10.1093/gpbjnl/qzad009

Ishii, T., Matsumoto, N., Tanaka, H., Eltayeb, A. E., & Tsujimoto, H. (2012). Evolution of subtelomeric and centromeric repetitive sequences in genus Pennisetum (Poaceae). Chromosome Science, 15(3+4), 53–59. 10.11352/scr.15.53

Kumar, S., Stecher, G., Li, M., Knyaz, C., & Tamura, K. (2018). MEGA X: Molecular Evolutionary Genetics Analysis across Computing Platforms. Molecular Biology and Evolution, 35(6), 1547–1549. 10.1093/molbev/msy096

Martel, E., Poncet, V., Lamy, F., Siljak-Yakovlev, S., Lejeune, B., & Sarr, A. (2004). Chromosome evolution of Pennisetum species (Poaceae): Implications of ITS phylogeny. Plant Systematics and Evolution, 249(3–4), 139–149. 10.1007/s00606-004-0191-6

Neumann, P., Novák, P., Hoštáková, N., & Macas, J. (2019). Systematic survey of plant LTR-retrotransposons elucidates phylogenetic relationships of their polyprotein domains and provides a reference for element classification. Mobile DNA, 10(1), 1. 10.1186/s13100-018-0144-1

Novák, P., Neumann, P., & Macas, J. (2020). Global analysis of repetitive DNA from unassembled sequence reads using RepeatExplorer2. Nature Protocols, 15(11), 3745– 3776. 10.1038/s41596-020-0400-y

Paudel, D., Kannan, B., Yang, X., Harris-Shultz, K., Thudi, M., Varshney, R. K., Altpeter, F., & Wang, J. (2018). Surveying the genome and constructing a high-density genetic map of napiergrass (Cenchrus purpureus Schumach). Scientific Reports, 8(1), 14419. 10.1038/s41598-018-32674-x

Rice, P., Longden, L., & Bleasby, A. (2000). EMBOSS: the European Molecular Biology Open Software Suite. Trends in Genetics : TIG, 16(6), 276–277. 10.1016/S0168-9525(00)02024-2

Sader, M., Vaio, M., Cauz-Santos, L. A., Dornelas, M. C., Vieira, M. L. C., Melo, N., & Pedrosa-Harand, A. (2021). Large vs small genomes in Passiflora: the influence of the mobilome and the satellitome. Planta, 253(4), 1–18. 10.1007/S00425-021-03598-0/FIGURES/5

Salson, M., Orjuela, J., Mariac, C., Zekraouï, L., Couderc, M., Arribat, S., Rodde, N., Faye, A., Kane, N. A., Tranchant-Dubreuil, C., Vigouroux, Y., & Berthouly-Salazar, C. (2023). An improved assembly of the pearl millet reference genome using Oxford Nanopore long reads and optical mapping. G3: Genes, Genomes, Genetics, 13(5). 10.1093/g3journal/jkad051

Sobrinho, F. D. S., Pereira, A. Vander, José, F., & Botrel, M. A. (2005). Avaliação agronômica de híbridos interespecíficos entre capim-elefante e milheto. Pesquisa Agropecuaria Brasileira, 40(9), 873–880.

Soltis, D. E., Soltis, P. S., Bennett, M. D., & Leitch, I. J. (2003). Evolution of genome size in the angiosperms. American Journal of Botany, 90(11), 1596–1603. 10.3732/ajb.90.11.1596

Techio, V. H., Davide, L. C., & Pereira, A. Vander. (2006). Meiosis in elephant grass (Pennisetum purpureum), pearl millet (Pennisetum glaucum) (Poaceae, Poales) and their interspecific hybrids. Genetics and Molecular Biology, 29(2), 353–362. 10.1590/S1415-47572006000200025

Varshney, R. K., Shi, C., Thudi, M., Mariac, C., Wallace, J., Qi, P., Zhang, H., Zhao, Y., Wang, X., Rathore, A., Srivastava, R. K., Chitikineni, A., Fan, G., Bajaj, P., Punnuri, S., Gupta, S. K., Wang, H., Jiang, Y., Couderc, M., … Xu, X. (2017). Pearl millet genome sequence provides a resource to improve agronomic traits in arid environments. Nature Biotechnology, 35(10), 969–976. 10.1038/nbt.3943

Wang, C., Yan, H., Li, J., & Zhou, S. (2018). Genome survey sequencing of purple elephant grass (Pennisetum purpureum Schum ‘Zise’) and identification of its SSR markers. Molecular Breeding, 39(194). 10.1007/s11032-018-0849-3

Yan, Q., Wu, F., Xu, P., Sun, Z., Li, J., Gao, L., Lu, L., Chen, D., Muktar, M., Jones, C., Yi, X., & Zhang, J. (2021). The elephant grass (Cenchrus purpureus) genome provides insights into anthocyanidin accumulation and fast growth. Molecular Ecology Resources, 21(2), 526–542. 10.1111/1755-0998.13271

Zhang, S., Xia, Z., Li, C., Wang, X., Lu, X., Zhang, W., Ma, H., Zhou, X., Zhang, W., Zhu, T., Liu, P., Liu, G., Wang, W., & Xia, T. (2022). Chromosome-scale genome assembly provides insights into speciation of allotetraploid and massive biomass accumulation of elephant grass (Pennisetum purpureum Schum.). Molecular Ecology Resources, 22(6), 2363–2378. 10.1111/1755-0998.13612

Zhang, S., Xia, Z., Zhang, W., Li, C., Wang, X., Lu, X., Zhao, X., Ma, H., Zhang, W., Zhou, X., Zhu, T., Liu, P., Liu, G., Yang, H., Arango, J., Peters, M., Wang, W., & Xia, T. (2020). Chromosome-Scale Genome Assembly Provides Insights into Speciation of Allotetraploid and Massive Biomass Accumulation of Elephant Grass (Pennisetum purpureum Schum.). 10.1101/2020.02.28.970749

