## Supplemental Material for "*Cenchrus purpureus* and *Cenchrus americanus* repeatome provide chromosomal markers to distinguish subgenomes"

Supplementary Table 1 – Satellite DNA Probes sequences and their respective source species and probe sequence.

| Satellite | Source species |
| --- | --- |
| CpaSat1A | *C. americanus*, *C. purpureus* |
| CpurSat5Y | *C. purpureus* |
| CpurSat7Y | *C. purpureus* |
| CpurSat9Z | *C. purpureus* |
| CameSat2B | *C. americanus* |

>CpaSat1A
CACAACTTTGGTACCCCGAAATAGTGCATTCAGGCCCGAAACACAAGTTTTGCATCTTTTT
ACGTGCCGAAGGTTTGCGAAATGCTCCGAAACACTCCCAAACATCATTTTGGGTCTAATGG
AGTAGAATGGATGCTT

>CpurSat5Y
GCCAAAGAAAATGACATGCGACTGGCTAAAGAACACTCTTCAAAACCATGTTGTCGTGCA
TTTCCAGCACAACCGGTGTC
GTTTTGATTGCCATCTAGCAGTAGAAACAATATACAAGGCCCTAATGCAGCCTTTACGCAT
GCTACTTTATGGGCCACAC
AATCTTTTGAGAGAAAATGTTTAAAAACCTGGTGAATAGCCACTTTTCTCACATTTTTTGG
CACAACCTTTGGAGGAACG
ATGCATTTTAGGGCCATCACCTTGAAGTTTCCATGGAAATAAGCCTGCAACAAGTTAATCC
AGTCCATTGTAGGCCCTTC
TATGCTGAGCATGTCA

>CpurSat7Y
CTGAAGATCTAAGCTGTGTTCCTTCCATCCTGAAAGTCCATAACTTTTCTGTTTTGAGCCCG
AATCGAGTGCGGTTTTTT
TTGTTAGATGCAGAATGCTCCTAGTTAGGTAATGCAAGGCGAATTTCCATGATTTGAGCAA
TTATTATTTTTATAAAAAT
AAAATCATGCCCACATGTGGAAAGATGCATCTCTTGGTCGTGGCTACGGTGCTGCTCTCGA
GAGGGCCTTGCTCGTACTA
GAATGGAAGGTTAACAACAGAAGTAATAGTCCGACAACACCGATACATGCCATATGCCAT
CCCAGGTGCATCGTTGCGAT
GTAAACCTTACGTTTCGGCCCGTTTCGTGGTGTATTAACACCTTCTATTGTG

>CpurSat9Y
CCATTTCGTTAATTTTTTGGTTACCTCAGGCCCCGGACAGTGACCAAAATGCACGGAAATT
CGCTAAAACATGTGTAAGA
GTGAAGAATACGGAGAATTTTCTTGAAAGTGGCAACTCTCCCAATCCCGGTTGACAAGTCT
AGGTGCCCTAGAGTCCATA
CAAGTAGTGGATACAATAGAACAAATATATGAGATGCAACACTTGCTGTTGGAGCACGAA
ACAAAGGCTATAACACACGG

TCGTAGAATTACTTTTGGTGAGGATTAGCCTTGTTATTTTTGTAAAAAAATAAATTGAGGT
GAACTCCTTAAAATTCATT
TTTCACGGTGTAGAATACGTCGTTATGTTTCCAACAAAAAAAAGAGCGCCAAAATCGGAC
TTGGGACGGAGAAGTTATGT
CATTTCAGGTAATGTGACCGTTGGTCCAATATTTGGGTAGTTTCTCTATGTAGATGCGGTCT
GCTACCCCCTGTGCATAC
CAGCAAAATTCATTTTGGTAACCACGAACAAGAATGTGAAAAAAATTG

>CameSat2B
TGGAGTGACGAAATGTGTCATTCTGGCTTCCGTTTGAGCAAGTAAATTTCTTCTAGGAAAG
CTAACGAAATGACCCGTAA
AAGGCCTGGAACGTGGACCCTATGGTCCAAAAATCAATCGAAACCTCATAAAATGTCATC
TACAAGTTTTTTGTGACAAT
CCGGGAGGTTTCCAAAAAATTGGCCCGGGTGCCGAAAACCGTGTGCTATAGCTCACGAAA
ATCGCCCGAAACGGACGTTT
TCGCGAAAAAATAAAATTCGAAACCGTCTGGCCCGGAAACCTACCCTGGTGGTGCTATGC
CTTAAACTATGGTCGGTACG
TGGAGAAGGGACCGT

Supplementary Figure 1. Transposable Elements quantification in *Cenchrus purpureus* and *Cenchrus americanus*

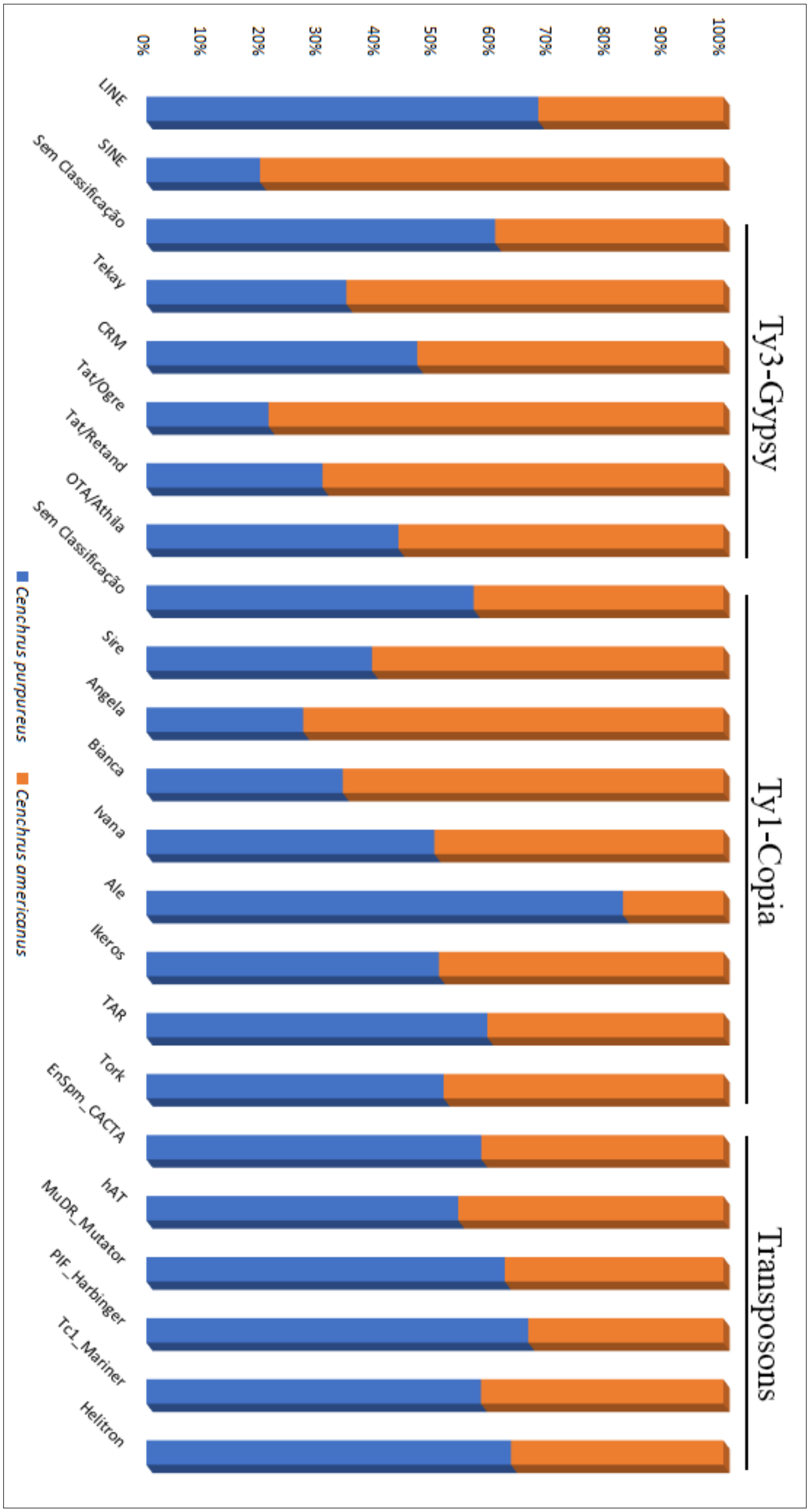
